# Discovering data-driven microbial growth models with symbolic regression

**DOI:** 10.1101/2025.06.02.657040

**Authors:** T. Anthony Sun, Dovydas Kičiatovas, Inga-Katariina Aapalampi, Teemu Kuosmanen, Teppo Hiltunen, Ville Mustonen

## Abstract

1. Connecting mathematical models with empirically measured microbial growth has remained challenging, as numerous competing models based on different theoretical approaches can fit observations. Therefore, we develop a method to automatically propose growth models from microbial data alone. We validate this approach using an available dataset of *E. coli* grown on known resources, and study sixteen species across various concentrations of a rich medium.
2. The inherently interpretable approach of symbolic regression infers explicit dynamical models directly from growth data. Using symbolic regression natively, does not favour biologically interpretable models, but we find cumulative population gain to be a more informative machine learning feature than population size.
3. Random Forest machine learning allows us to relate this finding to the approximation of a constant-rate per capita resource consumption. This suggests that the area under the growth curve (AUC) measured in routine experiments provides information on the effective resource dynamics governing microbial growth. Finally, we use theoretical insights to inform the symbolic regression algorithm and favour biologically interpretable models.
4. Overall, we found that balancing between data fit, parsimony and biological relevance favoured both the simplest, linear approximation, and models based on a Monod dynamics, with either one or two underlying resources. Therefore, our approach to read growth laws off of microbial batch cultures provides insights on data-driven modelling.

## 1 Introduction

Microbial growth is a key process in the ecology and evolution of microbes. For instance, understanding it quantitatively is essential for public health in the face of rising antibiotic resistances in human pathogens (Antimicrobial Resistance Collaborators, 2022; Wong, 2024), and for various biotechnology applications (Lässig et al., 2023; Yeoman et al., 2020). Yet, quantifying how microbial populations convert resources into biomass remains challenging, with most experiments relying on chemically complex media and short culturing times. So far, there has been no simple way to use batch culture data to infer how microbial populations grow, without relying on strong prior modelling assumptions. We know for certain that a growing microbial population uptakes resources from the environment and converts them to new biomass, and previous research has established that growth in batch cultures involve two intrinsic nonlinearities. On the one hand, the typical lag phase observed in growth experiments (Allen and Waclaw, 2018) implies that minimal models must include a time-dependent adjustment function to account for physiological changes occurring in bacterial cells encountering a fresh culture medium (Baranyi and Roberts, 1994; Huang, 2011; Ram et al., 2019a; Borse et al., 2024). On the other hand, the maximum per capita growth rate has a saturating response to an increase in resource concentration, demonstrated by Monod’s classical work on a single resource (Monod, 1949).

A wealth of theoretical work has proposed models for multi-resource growth (Zwietering et al., 1990; Baranyi and Roberts, 1994; Huang, 2011; De Jong et al., 2017; Allen and Waclaw, 2018; Letten, 2022; Held et al., 2024; Ghenu et al., 2024). Yet, quantitative analyses of empirical bacterial growth have often circumvented any explicit account of resource dynamics by using traditional models based on logistic growth (Ram et al., 2019a; Asnicar et al., 2023), which might misrepresent microbiological processes at work (Turchin, 2001; Balsa-Canto et al., 2019; Ram et al., 2019b; Picot et al., 2023; Hatton et al., 2024; Ishizawa et al., 2024). Our inability to fully use the biologically more appropriate resource-consumer models does not arise from a lack of theoretical tools (Cui et al., 2024), but from the fact that model complexity often far surpasses experimental evidence. This evidence typically covers observed population data but not resource dynamics, which creates a disconnect between observable phenotypes related to macroscopic growth (*e*.*g*., yield, effective growth rate, lag time) and underlying microscopic mechanisms (*e*.*g*., maximum growth rate, resource consumption) (Miguel Trabajo et al., 2024).

Indeed, several incompatible multi-resource models may fit experimental data equally well and produce, in effect, indistinguishable simulated observations (Breiman, 2001; Peleg and Corradini, 2011; Tang and Riley, 2021; Ghenu et al., 2024; Held and Manhart, 2024). This means that, in addition to the widespread question of parameter identifiability (Balsa-Canto et al., 2019), models are themselves not generally discoverable at realistic resolutions in microbiology. Even with an improved experimental resolution, we know that no algorithm can definitively find the generative model in arbitrarily complex cases (Richardson, 1969) nor in microbiological conditions (Shumaylov et al., 2025). The problem is that the various theoretical approaches to generate hypotheses themselves cannot be criticised on the basis of data: since traditional processes first assume a set of theoretical models, then gather empirical observations to fit the models before subsequent model selection, the initial choice of models itself is methodologically immune to data-based criticism. Thus, how the data could allow us to identify a true, generative model of microbial growth, is not generally a well-posed problem, outside the ideal case of perfect knowledge of all latent variables.

Therefore, we use a radically different approach of hypothesis generation, which is to accept working with approximations sufficient for specific use cases. This means asking which models suffice to reproduce the observed behaviour, to propose null-hypotheses *allowed* by empirical observations, irrespective of our theoretical priors, at first. In that sense, a good null-model trades off among data-drivenness, goodness of fit, biological interpretability and parsimony. In this data-driven approach, generating models as null-hypotheses should not primarily result from our assumptions but from finding patterns in the data, before model selection. Since theoretical input does not occur at the model proposal stage, it now becomes an essential part of model selection: after gathering empirical data, we should propose theoretical models in an unbiased way to fit data and, finally, filter model proposals through model selection, which now includes verifying that they make biological sense.

So far, an interpretable, yet automated, method to uncover dynamical growth models from experimental data has been missing. One interpretable alternative to black-box machine learning is symbolic regression, a method for the unbiased discovery of relationships between predictors and a dependent variable (Kronberger et al., 2025). By directly searching the space of mathematical expressions, it can fruitfully serve in hypothesis generation (Radwan et al., 2025). While computationally intensive, computing power advancements in recent decades have made symbolic regression viable and increasingly popular across a multitude of scientific disciplines, yielding impressive results in recovering and expanding known laws in physics (Course and Nair, 2023) and, in the context of microbial ecology, in reverse-engineering Lotka-Volterra and logistic-based models (Gaucel et al., 2014; Martin et al., 2018; Regalado, 2021; Hsin et al., 2023). At its basic usage, this methodology requires no assumptions or any other inputs than the data itself, but its true potential is in the ability to specify various constraints for the mathematical expressions by defining allowed mathematical operations and partial functional expressions. The latter constraint is extremely important for utilisation of the known mathematical laws and general domain knowledge, as showcased in this study. Symbolic regression is not an alternative to existing models, but an alternative to the approach of specifying candidate models *a priori*.

Here, we make minimal theoretical assumptions to better understand growth data obtained for fourteen bacterial species in Reasoner’s 2A (R2A) medium (figure 1). Since our ultimate goal is to propose models in arbitrary experimental settings with tractable or intractable resources, we use a complex medium where resources are not tightly controlled, to represent the general case, in addition to validating our approach using *E. coli* data obtained with known chemical resources (Held et al., 2024). The aim is to infer an effective resource dimensionality telling us how complex the medium is, from the point of view of a growing population of bacteria. We demonstrate that symbolic regression can reverse-engineer explicit dynamical models for the microscopic growth process. We find cumulative population gain to be a more informative machine learning feature than population size, then link this finding to the approximation of a constant-rate per capita resource consumption. We use theoretical insights to inform the symbolic regression algorithm and favour biologically interpretable models. The algorithm suggests that bacterial growth widely follows a Monod dynamics in our experiments, with either one or two effective underlying resources, even in our chemically complex medium. This is generally a slightly a better approximation than a linear-consumption model, but it remarkably applies to even a complex medium.

**Figure 1:**
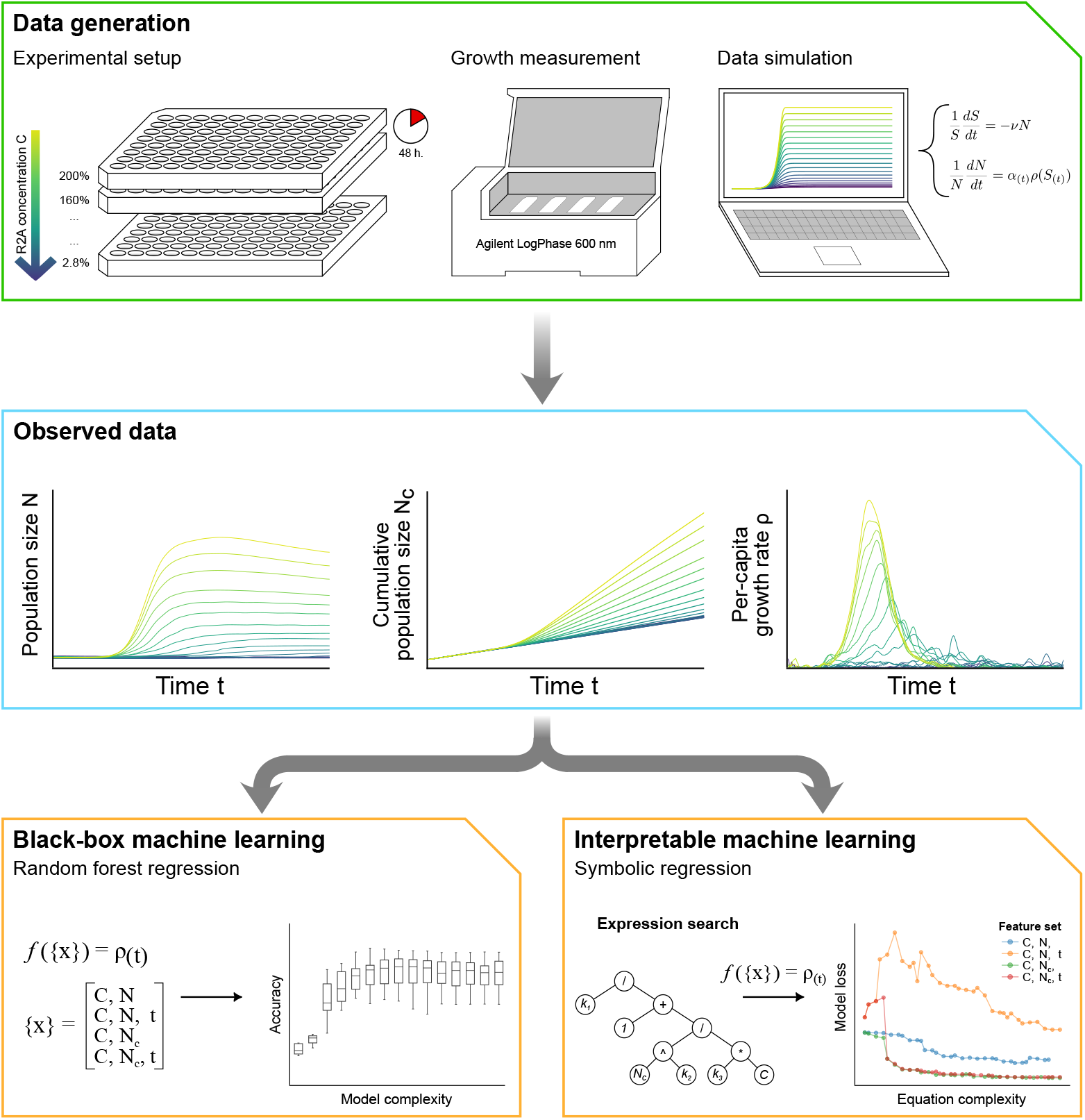
Workflow for the production and analysis of the datasets.

## 2 Materials and methods

### 2.1 Experimental methods for bacterial cultures

Our experiments involved 16 species from the HAMBI collection (Helsinki Institute of Life Science HiLIFE, Biodiversity Collections Research Infrastructure HUBCRI, Microbial Domain Biological Resource Centre HAMBI) summarised in Table S1. All were able to grow in Reasoner’s 2A medium and the following culture protocol yielded our data, summarised in Figure S1.

Solution M9 contained 11.28 g of M9 minimal medium salts (MP Biomedicals, lot `142662`) in 1.0 L of Milli-Q purified water. The culture medium (100 %) was prepared by dissolving then autoclaving 3.0 g of Reasoner’s 2A broth powder (Neogen, lot `UK208078/190`) into 1.0 L of solution M9. We prepared a range of 20 different concentrations *C* of culture medium, separated by an incremental factor 0.8, from 200 % to 2.88 %, by successive dilutions into autoclaved solution M9. Bacterial pre-cultures were run by inoculating each species into 6.0 mL of culture medium (100 %), then incubating for 48 h in aerobic conditions at 30 °C under agitation at 70 rpm, and finally diluted into autoclaved solution M9 by a factor 10^−4^.

Monocultures were run in four batch replicates by inoculating 110 µL of each culture medium concentration *C* with 20 µL of pre-culture onto a 96-well plate, then incubating for 48 h in aerobic conditions at 30 °C under agitation at 800 rpm. An Agilent BioTek LogPhase 600 device automatically recorded time and optical density at 600 nm every 10 min as a proxy to population size, yielding 16 species × 4 biological replicates × 20 concentrations = 1280 experimental growth curves, each comprising 289 time points. We considered 14 among the 16 species because two did not reach stationary phase. We derived all other observations numerically.

### 2.2 Theoretical methods and models

Consider a bacterial population of size *N* growing in a substrate of concentration *S*. The variables *S* and *N* are time-dependent. To account for the lag period observed in microbiological batch cultures, minimal models of bacterial growth assume a time-dependent physiological adjustment *α* in their population dynamics (Zwietering et al., 1990; Baranyi and Roberts, 1994; Ghenu et al., 2024):

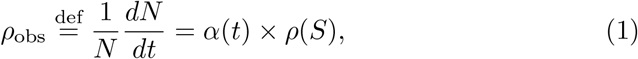

where *ρ* is the time-independent and potentially resource-dependent growth function. How physiological adjustment delays bacterial cells’ growth dynamics is usually represented by similar mathematical forms (Baranyi and Roberts, 1994; Huang, 2011; Lo Grasso et al., 2023; Borse et al., 2024; Ghenu et al., 2024). From fits to real data, we know that just two phenomenological parameters suffice to capture the adjustment function (Ram et al., 2019a; Borse et al., 2024), namely the initial physiological state of the cell (population) *q* (dimensionless) and the rate of change *m* (per unit time) in adjusting its physiology to the present environmental state:

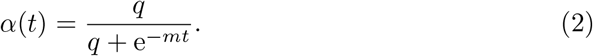

Monod (1949) showed that the maximum *ρ*_max_ of the empirical per capita growth rate *ρ*_obs_ was an increasing then saturating function of the resource concentration. The demonstration concerns only the maximum per capita growth rate *ρ*_max_(*S*) in a bacterial growth experiment (Monod, 1949; Angaroni et al., 2025). However, this type II functional response pattern (Holling, 1959a) is usually modelled using a hyperbolic function (Michaelis and Menten, 1913; Monod, 1949; Holling, 1959b) and considered a function of time within a dynamical model (Miguel Trabajo et al., 2024):

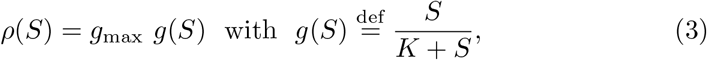

where *K* is the substrate concentration inducing half-maximum response. Without further assumptions, this functional form cannot model multi-resource growth (Monod, 1949), which may be complicated by non-mutually-exclusive resources (Perrin et al., 2020), colimitation (Saito et al., 2008; Held et al., 2024), and sequential preference (Kremling et al., 2018). As it is unclear how several resources (*S*_0_, …, *S*_*s*_) together impact bacterial growth, multi-resource growth models combine each resource’s Monod contribution to the total growth in various ways, notably with multiplicative or additive effects (Saito et al., 2008; Letten, 2022; Held and Manhart, 2024; Held et al., 2024):

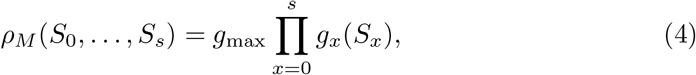

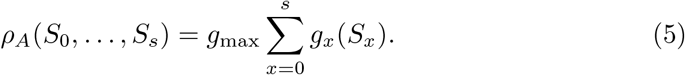

The multiplicative-effect multi-resource model combines Eq. (1) and Eq. (4). It assumes that growth requires all resources, which are thus biologically not substitutable. The additive-effect multi-resource model combines Eq. (1) and Eq. (5). It assumes that any resource allows for some growth, which implies sub-stitutable resources. A common framework can unify several formulations (Tang and Riley, 2021), but at the cost of a combinatorial explosion of theoretical parameters, as the number of resources increases. Overall, experimental data provides no definitive, general support for either model over the other (Tang and Riley, 2021; Held and Manhart, 2024), so that we have no prior hypothesis in our experimental setting involving several species and a rich medium.

### 2.3 Numerical methods

Each dataset consisted of a range of bacterial growth curves at different initial medium concentrations *C* for a given species. We used Python (3.10.14) for analysis, NumPy (1.26.4) and Pandas (2.2.2) for numerical protocols, Matplotlib (3.8.4) for visualisations, Scikit-learn (1.4.2) for Random Forest methods, PySR (1.3.1) for symbolic regression (Cranmer, 2023). Code is available at: https://github.com/anthony-sun/SR-Monod. Since the experimental batch replicates were extremely similar, we averaged population sizes across the four replicates, for each time point. This has no impact on the study in practice, due to a very low coefficient of variation (median interreplicate-CV: 1.3 %), but greatly facilitates computation. The supplementary information details the protocol used for simulated data and expands on the symbolic regression methods.

For symbolic regression, the aim was to find explicit mathematical growth models which we could then interpret, in light of biological knowledge, as revealing latent resources. Alternative frameworks exist, such as general additive modelling (Wood, 2025), with specific constraints on how variables may combine. We split each dataset into a training set (50 %) and a testing set (50 %) comprising every other growth curve along the range of medium concentrations. This ratio allowed for a sufficient testing dataset keeping the time series structure of whole growth curves, and avoided artefacts which could result from an uneven spacing of the training/testing medium concentrations. In any case, it was not possible for this unusual ratio to unduly inflate the performance of the symbolic regressor, much to the contrary. For a given complexity bound on output expressions, the algorithm optimised data fits based on mean squared-error using the default method (with crossover and mutation) and parameters in reference (Cranmer, 2023). More complex functional forms resulted from randomly combining feature variables and constants using: addition (+), multiplication (×), division (*/*), and constant power (^^*k*^, *k >* 0). We imposed biology grounded restrictions against negative constants and non-integer powers of the relative medium concentration *C*. To facilitate numerical computations, we normalised each dataset’s cumulative population gain *N*_*c*_ by its maximum value, and we used the exponential negative scaled time *U* = e^−*t*^ and the exponential negative cumulative population gain 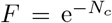 as features. Finally, we used expression templates constraining the search for mathematical expressions as detailed in the results.

For Random Forest regression, the aim was to suggest which observed variables would best predict growth rate and thus better reflected a latent resource dynamics. We split each dataset into a training set (75 %) and a testing set (25 %) comprising every fourth medium concentration along the range of medium concentrations. Since we were not interested in evaluating the Random Forest methods *per se*, but to see how various feature sets enable prediction, it was not critical to avoid artefacts which could result from an uneven spacing of the training/testing medium concentrations. The predicted response variable was the per capita growth rate *ρ*_obs_. Feature variables included: one population size feature, namely *N* or *N*_*c*_, the relative initial medium concentration *C*, the time *t*. We used all default settings of the Scikit-learn package, namely 100 estimators, the coefficient of determination *R*^2^ as a performance score, and no cap on the maximum tree depth by default, unless otherwise specified.

## 3 Results

### 3.1 Unconstrained symbolic regression predicts growth

Our hypothesis was that growth curves in a range of medium concentrations should allow for the inference of dynamical growth models. We demonstrated it with a symbolic regression algorithm. Symbolic regression consists of optimising a functional form predicting a response variable under constraints on formula complexity, thus providing explicit mathematical models with little theoretical bias (Cranmer, 2023). We used this tool to explore which key observables are more appropriate for a blind model search.

We found that models relying on cumulative population gain *N*_*c*_ systematically outperformed models relying on population size *N* (figure 2, {*C, N*} and {*C, N*_*c*_}) in terms of fitting performance. Feature set {*C, N*_*c*_} yielded the best overall performance (figure 2), especially with more complex models. Thus, our naïve symbolic regression algorithm showcases the potential for *N*_*c*_ to inform us on microbial growth resources while providing mathematically explicit models. Yet, it could not yield substantial biological insight beyond the importance of *N*_*c*_, because the models that it inferred optimised only their fitting performance and did not generally reflect basic theoretical expectations from the literature.

**Figure 2:**
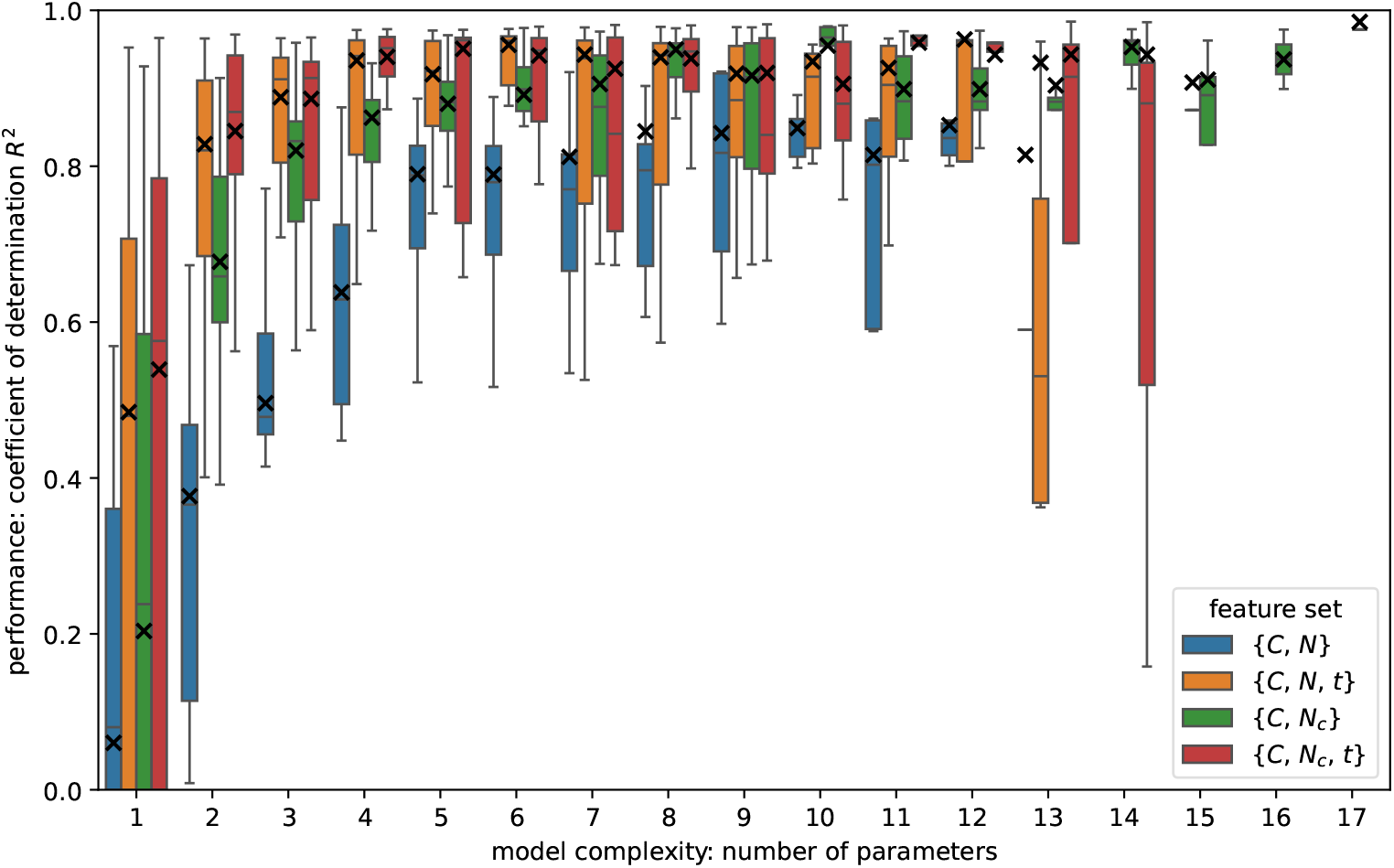
Performance of the unconstrained symbolic regressor in predicting the per capita growth rate from different feature sets in experimental data, given by the coefficient of determination *R*^2^. Crosses show the median train score across species. The absence of highly complex models when using certain feature sets means that the symbolic regressor did not propose any highly complex model. The cumulative population gain predictor *N*_*c*_ outperforms the population size predictor *N* as well as its combination with time *t*.

### 3.2 Cumulative population gain allows for black-box prediction of growth

Before addressing the issue of biological relevance, we asked why the cumulative population gain feature *N*_*c*_ yielded better results than population size *N* alone. We found that the observed behaviour was at least consistent with the theoretical approximation that resource consumption is a constant-per-capita-rate process. Indeed, the law of mass action assumes that a bacterial population of size *N* growing in a substrate of concentration *S* uptakes resources at a constant rate *ν* relative to its size:

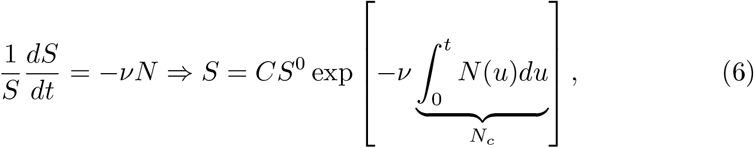

where *C* is the relative initial concentration of the growth medium compared to a reference, and *S*^0^ = *S*_(*C*=1,*t*=0)_ is the resource density in the reference concentration. This constant-rate model for per capita resource consumption readily extends to several independent resources and naturally gives a paramount role to the cumulative population gain *N*_*c*_. In laboratory terms, the cumulative population gain *N*_*c*_ corresponds to the area under the (growth) curve (AUC) sometimes reported in microbiology.

This theoretical expectation yielded a testable implication, independent from symbolic regression: models relying on the traditional, instantaneous population size *N* should perform worse than models relying on the cumulative population gain *N*_*c*_, but could be supplemented by providing time data *t* to approximate the integral in Eq. (6). To test that independently from symbolic regression, we compared the performance of a black-box Random Forest regressor using either *N* or *N*_*c*_. Our aim was not to optimise the performance of the machine learning algorithm but to compare the use of different feature sets.

Effectively, using *N*_*c*_ instead of *N* improved the performance of a Random Forest regressor in predicting per capita growth rate *ρ*_obs_ (figure 3, {*C, N*} and {*C, N*_*c*_}; two-sided correlated t-test on the models with no maximal tree depth: *p* = 0.0011). Feature *N*_*c*_ effectively captured the impact of both *N* and *t* since feature sets {*C, N, t*}, {*C, N*_*c*_}, and {*C, N*_*c*_, *t*} achieved a similar improvement. They also yielded simpler decision trees: while relying on *N* led to a median maximum tree depth of 35, relying on *N*_*c*_ instead reduced it to 25 (table 1). When capping the maximum depth of all decision trees from the start, all three yielded a systematic improvement of the Random Forest regression compared to feature set {*C, N*} (figure 3).

**Table 1:** Unrestricted complexity of a Random Forest regressor using 100 trees to predict the per capita growth rate: median across 14 experimental datasets, each comprising 5780 points and corresponding to one species.

| feature set | train score | test score | median maximum tree depth (SD) |
| --- | --- | --- | --- |
| $\{C, N\}$ | 98.1 % | 72.0 % | 35 (5) |
| $\{C, N, t\}$ | 99.9 % | 91.3 % | 27 (4) |
| $\{C, N_c\}$ | 99.9 % | 91.6 % | 25 (3) |
| $\{C, N_c, t\}$ | 99.9 % | 93.0 % | 27 (3) |

**Figure 3:**
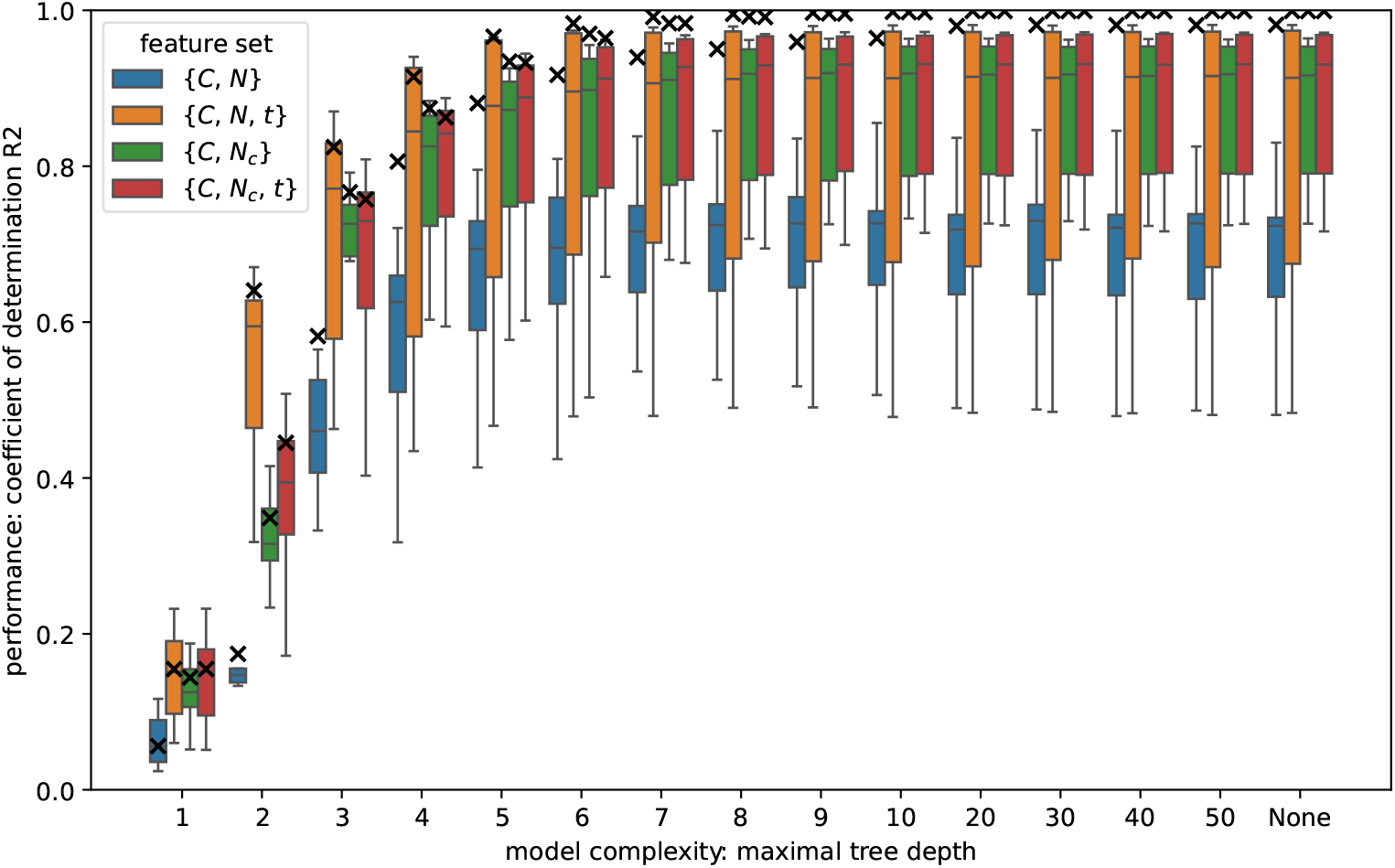
Performance of a black-box Random Forest regressor in predicting the per capita growth rate from different feature sets in experimental data, given by the coefficient of determination *R*^2^. Crosses show the median train score across species. The cumulative population gain predictor *N*_*c*_ outperforms the population size predictor *N*.

When relying on population size *N* to predict the per capita growth rate *ρ*_obs_, the Random Forest regressor showed systematic overfitting, with a gap between train and test scores. Models relying instead on cumulative population gain *N*_*c*_ reduced this overfit.

The feature sets {*C, N, t*}, {*C, N*_*c*_} and {*C, N*_*c*_, *t*} roughly yielded a similar improvement. The theoretical formulation of Eq. (6), implying a map from resource *S* onto the unidimensional space *N*_*c*_, suggested that the algorithms could use *N* and *t* together to computationally approximate *N*_*c*_. In any case, *N*_*c*_ effectively offered a parsimonious, unidimensional approximation of the bidimensional spaces (*N, t*) and (*N*_*c*_, *t*).

Our data confirmed that, compared to population size *N*, cumulative population gain *N*_*c*_ improved the performance of a Random Forest black-box, including by reducing the overfitting. This cannot *prove* that resource consumption has a constant per capita rate but showed that this was a reasonable approximation, consistent with the data in our experiments.

### 3.3 Constrained symbolic regression infers biological growth model

Next, we hypothesised that a symbolic regression algorithm could infer a biologically interpretable growth function, when guided by theory-based expectations. Thus, we ran three rounds of symbolic regression: one with minimal biological assumptions (unconstrained as previously), and two informed by supplementary theoretical assumptions, using the following expression templates:

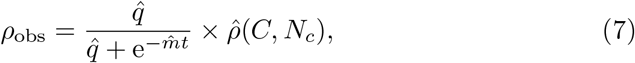

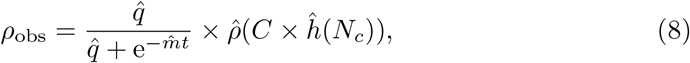

where the algorithm looked for the undefined parameters 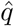 and 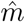, and for the undefined functions 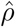 and *ĥ*. Note that those templates effectively mean using feature set {*C, N*_*c*_, *t*}. Template (7) assumed Eq. (1) and Eq. (2) by imposing the search for adjustment parameters *q* and *m*. Template (8) inspired by Eq. (6) additionally assumed that the growth’s resource-dependency is proportional to the medium’s relative concentration *C*.

We focused on how symbolic regression could inform microbial growth in two use cases corresponding to different research questions. In the first use case, our approach determined the biological substitutability or non-substitutability of known resources. In the second use case, it inferred the effective dimensionality of the resource space, as perceived by a growing bacterial population, when the exact limiting resources were not known.

#### 3.3.1 Inferring non-substitutability of known resources

The first use case concerns simulations and empirical data in which chemical resources are known and varied independently from one another in the growth medium. This experimental setting allows for a modelling investigation of whether resources are substitutable or not. We simulated growth curves using a dynamical Monod model with either a non-substitutable or substitutable bidimensional resource, combining respectively Eq. (1), Eq. (2), Eq. (3), Eq. (4) and Eq. (6), or Eq. (1), Eq. (2), Eq. (3), Eq. (5) and Eq. (6). Using data obtained under independently varying resource conditions, the symbolic regressor was able to correctly learn the substitutability of the provided resources (figure S4).

We repeated those observations with available batch culture data where *E. coli* was grown on glucose as a carbon source and ammonium as a nitrogen source (Held et al., 2024). Among the best-fitting, biologically interpretable models proposed by the algorithms corresponded to a multiplicative, bidimensional, dynamical Monod formulation (figure 4). This multiplicative model biologically implied that the two inferred resources were not substitutable, in agreement with the fact that glucose was the only carbon source, and ammonium, the only nitrogen source. Moreover, the fact that the one-resource models proposed relied on glucose concentration rather than on nitrogen concentration, aligns with the key impact of the carbon source on the growth dynamics.

**Figure 4:**
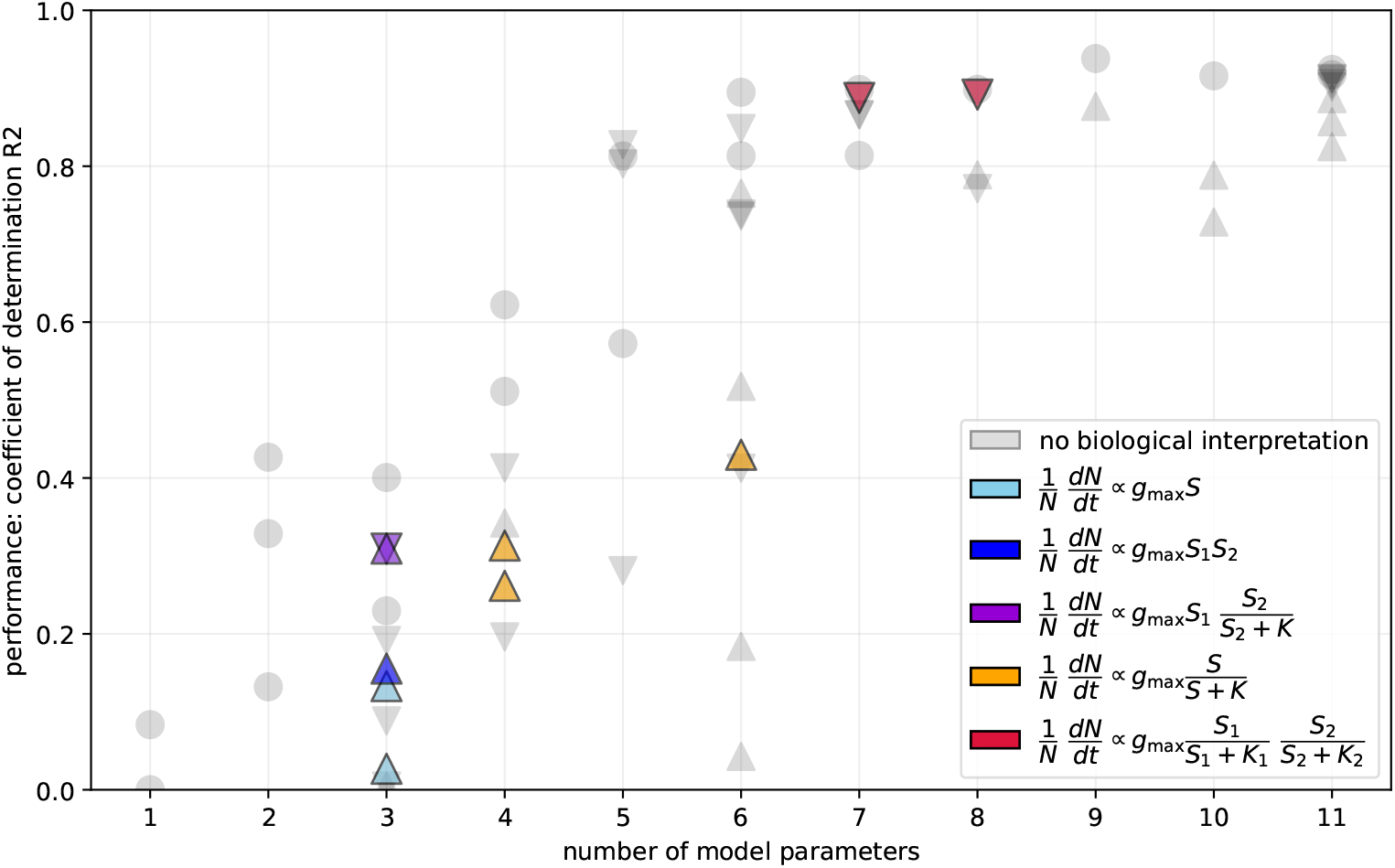
Fitting performance of models proposed by the symbolic regressor from growth data obtained with batch cultures of *E. coli* on known resources: glucose and ammonium, which are non-substitutable. Circle: no template; triangle pointing down: Template (7); triangle pointing up: Template (8).

This provides for a validation of our approach through simulations and experimental data. Namely, in the ideal case where resources are known, our approach is able to inform us on their biologically relevant properties, such as their substitutability.

#### 3.3.2 Inferring effective dimensionality of unknown resources

The second use case concerns the more general situation where resources are not tightly controlled, not precisely known or not chemically well-defined. There, the question is rather how a given microbial species perceives a given medium: what is the effective dimensionality of this medium as reflected by the growth dynamics? Experimental datasets were obtained in R2A, a rich culture medium containing yeast extract, thus yielding no expectations regarding resource limitations. The algorithms generated 1068 models compatible with the data. A data-driven approach cannot prove nor disprove any mathematical expression: in simulated data for example, it may or may not propose the generative model (Supplementary information) with no noticeable difference in fitting abilities or other performance metrics. This is an expected consequence of the known lack of model discoverability (Balsa-Canto et al., 2019; Tang and Riley, 2021; Shumaylov et al., 2025). Instead, a rigorous interpretation is that proposed models are sufficient to approximate empirical observations. To avoid focusing on well-fitting models with no biological interpretability, we manually annotated models to pick those which were compatible with theoretical expectations (Supplementary information). Overall, we found few of the proposed models to be interpretable in light of the literature in microbiology (figure 5).

**Figure 5:**
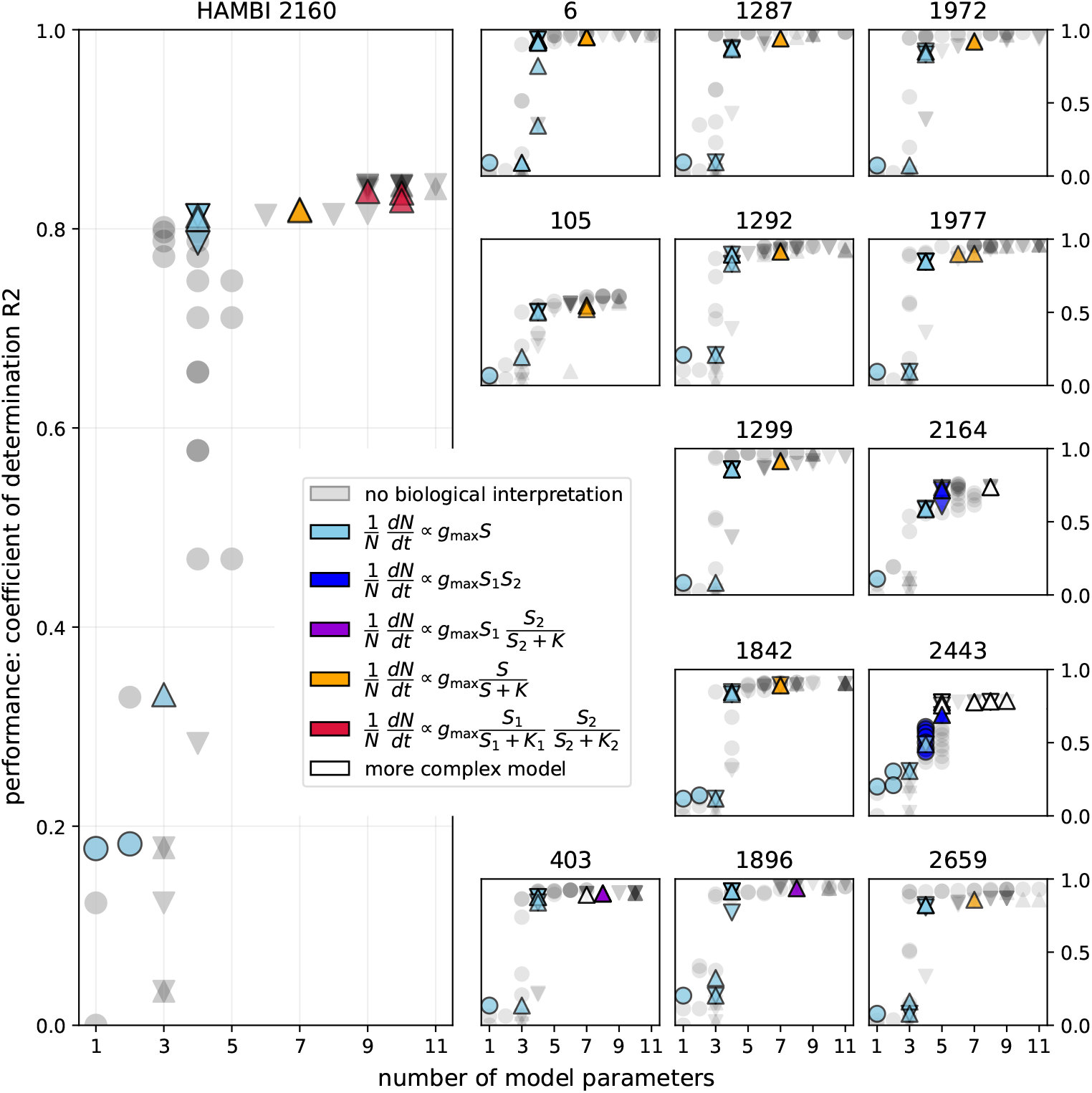
Fitting performance of models proposed by the symbolic regressor for R2A data with unknown resource dimensionality. Circle: no template; triangle pointing down: Template (7); triangle pointing up: Template (8).

Table 2 summarises the best-fitting models that remained biologically interpretable in experiments. In many species, the algorithm initially proposed a linear model, where the resource-dependent growth *ρ* was a linear function of a unidimensional, latent environmental resource *S*. This remained the best interpretable model for one of 16 experimental species in R2A medium, suggesting that experimental conditions did not cover the saturating part of Monod’s formulation. In all other species indeed, we retrieved a dynamical formulation of the Monod model corresponding to Eq. (1) and Eq. (6), which slightly improved fits. This also implied an effectively unidimensional resource space where the per capita growth rate was a saturating function of a single environmental resource. In four cases, the algorithm further found that the observed growth was better explained by an effectively multidimensional resource, with constant-per-capita-rate or Monod-like components.

**Table 2:** Formula, interpretation and inferred resource dimensionality of the best-fitting, biologically interpretable growth models for *ρ* in Eq. (1) and Eq. (3).

| model for $\rho(S)$ | interpretation | res. dim. | number of experimental species |
| --- | --- | --- | --- |
| $g_{\max}S$ | linear | 1 | 1 sp.: 2443 |
| $g_{\max} \frac{S}{K+S}$ | Monod | 1 | 9 sp. |
| $g_{\max}S_1 \times \frac{S_2}{K+S_2}$ | linear × Monod | 2 | 2 sp.: 403, 1896 |
| $g_{\max} \frac{S_1}{K_1+S_1} \times \frac{S_2}{K_2+S_2}$ | Monod × Monod | 2 | 1 sp.: 2160 |
| more complex | linear and Monod | $\geq 2$ | 1 sp.: 2164 |

However, the fitting improvement brought about by more complex models was often limited (figure 5). This means that linear models are good first approximations in many cases. Modellers may improve fitting performance by considering instead Monod’s model, which has a limited cost in terms of parameters and complexity. Whether the fitting improvement is worth the increased complexity will depend on the specific objective at hand, especially when considering multidimensional Monod models.

## 4 Discussion

### 4.1 Biological implications

In this study, we used symbolic regression to learn explicit growth models from population size data, without relying on prior theory. While fits to the data were excellent, an unconstrained use of the method did not ensure biological relevance, thus emphasising the need for a biologically bespoke usage of symbolic regression. However, our approach suggested that cumulative population gain *N*_*c*_ provides a powerful and simple way to indirectly observe the latent resource space experienced by growing microbial populations. We confirmed this using Random forest machine learning and linked it with theoretical expectations resulting from the law of mass action expressed by Eq. (6).

Finally, we combined symbolic regression with theoretical insights based on the literature, namely: biological constants are positive; expressions combine through addition, multiplication, division and power; biological mechanisms involve only integer powers of the medium concentration *C*; the law of mass action of Eq. (6) applies, *i*.*e*., *N*_*c*_ is an important state variable; the effective per capita growth rate follows Eq. (1) as the product of an adjustment function and a resource-dependent part. We validated our approach using an available dataset in which resources were known (Held et al., 2024) before applying it to our experimental data in R2A. We found that the dynamical resource-consumer model based on Monod’s formulation in Eq. (3), which we never assumed, best trades off among fit to data, symbolic simplicity, and biological interpretability, which extends previous non-dynamical findings (Angaroni et al., 2025). Our experimental observations effectively follow the approximation, formulated by Eq. (1) and Eq. (6), of a dynamical Monod growth model with latent resources consumed at a constant per capita rate. Overall, they highlight the potential of symbolic regression as a data-driven hypothesis generation method while emphasising the need to integrate existing theory and biological knowledge when interpreting its outputs.

Our approach also provides an access to the effective resource dimensionality experienced by a growing microbial population. The effective resource dimensionality represents the minimal number of effective, independent resources theoretically required to reproduce the observed population dynamics. In most cases, the growth dynamics can be approximated as relying on a unidimensional resource (Table 2) (Monod, 1949), such as a linear combination of the concentration of one or several compounds present in the medium. Even in our chemically complex medium, the number of effective resources remained remarkably low. The exact link between data-driven models and mechanistic resource dynamics is an excellent follow-up question raising several interpretative difficulties. For instance, if a complex medium can be approximated as relying on a unidimensional resource, which we may interpret as limiting, it does not follow that a single chemical acts as the key limiting resource throughout the growth experiment. Moreover, the fact that two species each sees the same medium as a unidimensional resource space, does not imply that they see the same linear combination of chemically defined resources, although this provides a useful null-hypothesis. Additionally, consider a chemically defined medium with a single limiting resource 1 yielding model *A* with resource 1^*′*^. If enriched with a second chemically defined resource 2, it could be approximated by a new model *B* with resources 1^*′*^ and 2^*′*^. There is no guarantee that model *B* reduces to model *A* when one of its resources is fixed.

The key point is that modelled effective resources are defined purely from their impact on the growth dynamics and, as is the case with all machine learning methods, reflect the data provided. Therefore, a strict interpretation of symbolic regression outputs cannot extend to mean that we reveal chemical resources. Departing from a chemical view allows us to translate population dynamics into effective ecological niches, which can be generally disconnected from chemical resources (Meszéna et al., 2006). Among multidimensional resource models, multiplicative effects suggest that latent resources are non-substitutable (Held and Manhart, 2024), whereas additive effects suggest that they are substitutable. Thus, our approach connects microbiology experiments with a rich literature on multidimensional niches (Hutchinson, 1961; MacArthur and Levins, 1967; Posfai et al., 2017) and their overlap, which determine species coexistence in communities (Chesson, 2018; Badali and Zilman, 2020; Pastore et al., 2021; Rozas Garcia et al., 2023). Future research could modulate the concentration of single chemical resources in a culture medium, to allow for a high-definition view of the growth process in a complex environment, and discuss the interpretation of symbolic regression outputs in light of more chemical evidence (Takano et al., 2023).

### 4.2 Methodological impact of symbolic regression in microbial ecology

Despite its success in physics (Course and Nair, 2023), symbolic regression has mainly been constrained to search logistic-based models in microbiology (Gaucel et al., 2014; Martin et al., 2018; Regalado, 2021; Hsin et al., 2023), thus overlooking two limitations. First, such models fail (Turchin, 2001; Balsa-Canto et al., 2019; Ram et al., 2019b; Picot et al., 2023; Hatton et al., 2024; Ishizawa et al., 2024) to account for key resource dynamics mediating the interactions (Meroz et al., 2024; Miguel Trabajo et al., 2024) that explain how bacterial communities assemble (Coyte et al., 2021; Lee et al., 2023; Ho et al., 2024), grow, organise and change (Pacheco et al., 2021), through competition and resource utilisation traits (Litchman et al., 2015; Ho et al., 2022). Secondly, assuming a functional form *a priori* leaves the plethora (Zwietering et al., 1990; Baranyi and Roberts, 1994; Huang, 2011; De Jong et al., 2017; Allen and Waclaw, 2018; Letten, 2022; Held et al., 2024; Ghenu et al., 2024) of growth functions proposed in the literature unexploited. Rather than developing more and more complex theory, it is essential to better connect empirical data on microbial growth with available models offering easy computation and best understanding.

We demonstrated how to combine pre-existing knowledge with the potential for symbolic regression to automatically infer an effective growth function from data, with little to no prior constraints on its functional form. Since symbolic regression does not provide automatic scientific interpretation, it is useful in generating, but not testing, hypotheses (Radwan et al., 2025). Clearly, it should not be confused for a tool to recover arbitrary generative models. Instead, symbolic regression constructs models that are sufficient to reproduce observations. This is independent from generative processes due to well-known parametric identifiability and model discoverability caveats (Ram et al., 2019b; Tang and Riley, 2021; Held and Manhart, 2024; Shumaylov et al., 2025). Notably, our proposed models involve exclusively observable variables. Only scientific interpretation in light of pre-existing knowledge expressed by Eq. (6) allows for a translation into a latent resource space. Indeed, symbolic regression outcomes cannot reject alternative hypotheses, which may still receive other empirical support, *e*.*g*., through knowledge about specific metabolites involved. Thus, future research should further investigate the link between effective resources inferred by symbolic regression and specific chemicals. A general rule-of-thumb is that assuming more resources than retrieved by symbolic regression is not necessary from the viewpoint of microbial growth.

The bottom line is that all biologically interpretable models proposed by symbolic regression could capture the observed behaviours and act as null hypotheses. Model proposals must be discussed based on goodness of fit, especially when comparing them, and in light of a specific research question at hand. Indeed, symbolic regression does not suffice to complete the model selection process, which is distributed between constraints on model search before symbolic regression, fit to data during symbolic regression, biological interpretability after symbolic regression, and balancing fits with model complexity. The specific coarse-graining level that researchers should apply to symbolic regression outputs depends on their particular use case. For several of our species indeed, the fitting improvement brought about by considering several resources is small, compared to a unidimensional resource. This means that researchers interested in a rough approximation can generally assume a unidimensional dynamical Monod model, while researchers interested in resource consumption of their species in a particular medium may focus on more complex models. In the extreme case, fitting a linear-consumption model remains the easiest approach, due to a low number of parameters, and often yields roughly acceptable fits. However, symbolic regression sets a null-hypothesis upper bound on resource dimensionality.

## 5 Conclusions

Our study demonstrates that symbolic regression offers a promising, empirically viable, and scalable way to analyse routine experiments in microbiology to infer null-models for growth and effective resource dimensionalities. Compared to the traditional method of fitting a posited family of models to data, our approach does not predefine candidate models. Instead, we build a cohort of candidate models based on the data first, then assess their theoretical and biological relevance. When resources were known and varied independently, the symbolic regressor was able to inform their metabolic interplay. When resources were not known, it was able to suggest an effective number of resources perceived by each species. We found that it is reasonable to assume a constant-rate per capita resource consumption and, based on this assumption, we expect the dynamical Monod model to capture the growth behaviour of most species in batch monocultures. Microbiologists can now assess this assumption using experimentally measured growth curves, and knowledge of resource consumption dynamics could inform our predictions and control of microbial community interactions.

## Acknowledgements

The authors wish to thank Frédéric Guillaume for comments on an earlier version of the manuscript, and CSC — IT Center for Science, Finland, for computational resources. This work for was in part supported by Research Council of Finland (345829, 346128, and 364234) to VM and TH.

## Notes

### Competing Interest Statement

The authors have declared no competing interest.

### Summary of Updates

Expand Introduction, Results (3.3) and Discussion. New Fig. 4, improved Fig. 5. Supplemental information updated.

