## Supplementary material for "Discovering data-driven microbial growth models with symbolic regression": SI

### Supplementary information

### 1 Overview of the experimental data

*Table 1: Bacterial species used in the experiments, and their HAMBI code. The last two did not generally reach stationary phase and were not considered for further analyses.*

| HAMBI | identification |
|---|---|
| 6 | *Pseudomonas putida* |
| 105 | *Agrobacterium tumefaciens* |
| 403 | *Comamonas testosteroni* |
| 1287 | *Citrobacter koseri* |
| 1292 | *Morganella morganii* |
| 1299 | *Kluyvera intermedia* |
| 1842 | *Sphingobium yanoikuyae* |
| 1896 | *Sphingobacterium spiritivorum* |
| 1972 | *Aeromonas caviae* |
| 1977 | *Pseudomonas chlororaphis* |
| 2160 | *Bordetella avium* |
| 2164 | *Cupriavidus oxalaticus* |
| 2443 | *Paracoccus denitrificans* |
| 2659 | *Stenotrophomonas maltophilia* |
| (3031) | *(Niabella yanshanensis)* |
| (3237) | *(Microvirga lotononidis sp. Nov.)* |

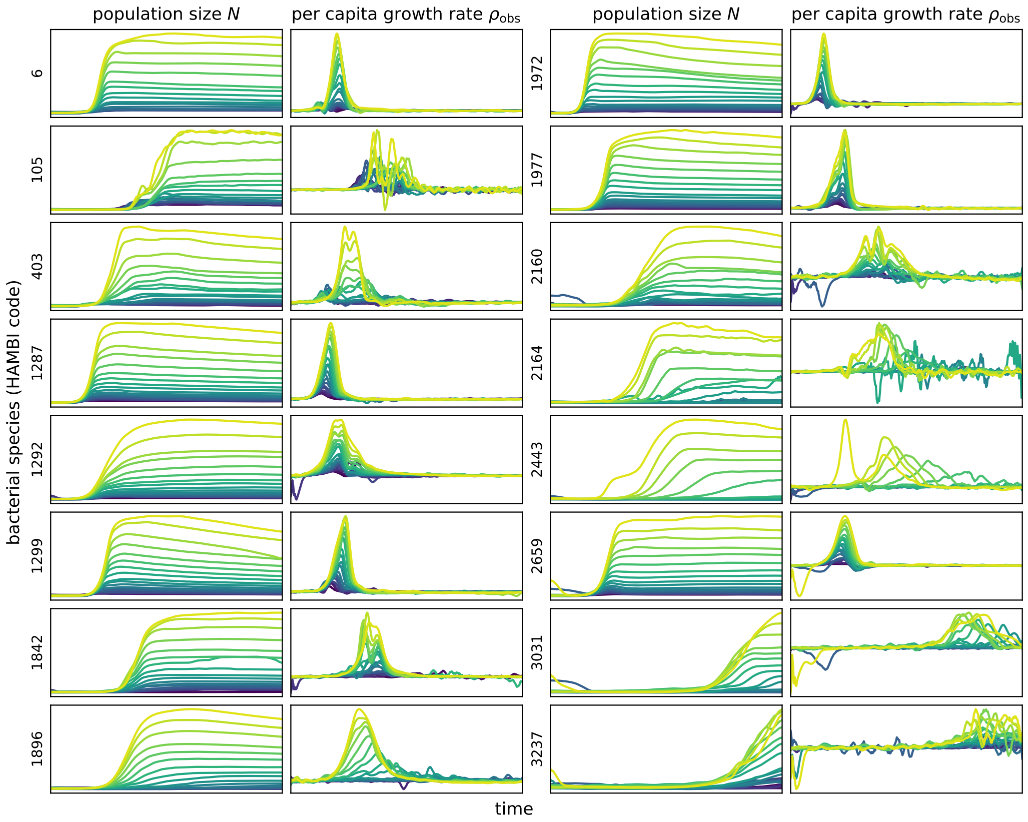

*Figure 1: Overview of the experimental data, showing the optical density of bacterial cultures and the per capita growth rate as a function of time.*

### 2 Random Forest regression

#### 2.1 Simulation for the Random Forest

We simulated three categories of models for comparison with the experimental dataset (labelled `Rs0`):

- the multiplicative-effect model `M` presented in the main text methods, combining Eq. (1), Eq. (3), Eq. (4) and Eq. (6), with resource dimensions between 1 and 5 (`Ms1` through `Ms5`), adjusted following Eq. (2);
- the additive-effect model `A` presented in the main text methods, combining Eq. (1), Eq. (3), Eq. (5) and Eq. (6), with resource dimensions between 1 and 5 (`As1` through `As5`), adjusted following Eq. (2);

Simulated species correspond to different sets of parameters, which were randomly generated in the orders of magnitude encountered in experimental datasets. Each dimension of the initial resource vector was set to 1; and the initial population size, to 0.1. We used the Runge-Kutta 4(5) method to integrate over the time span $[0, 50]$ with a time step of 1/6, matching the timing (h) of experimental samples. Cumulative population gain $N_c$ was computed using the trapezoidal rule.

For simulations, parameter ranges were randomly generated in the orders of magnitude encountered in experimental datasets. Each dimension of the initial resource vector was set to 1; and the initial population size, to 0.1. We used the Runge-Kutta 4(5) method to integrate over the time span $[0, 50]$ with a time step of 1/6, matching the timing (h) of experimental samples. Cumulative population gain $N_c$ was computed using the trapezoidal rule.

#### 2.2 Performance in simulated data

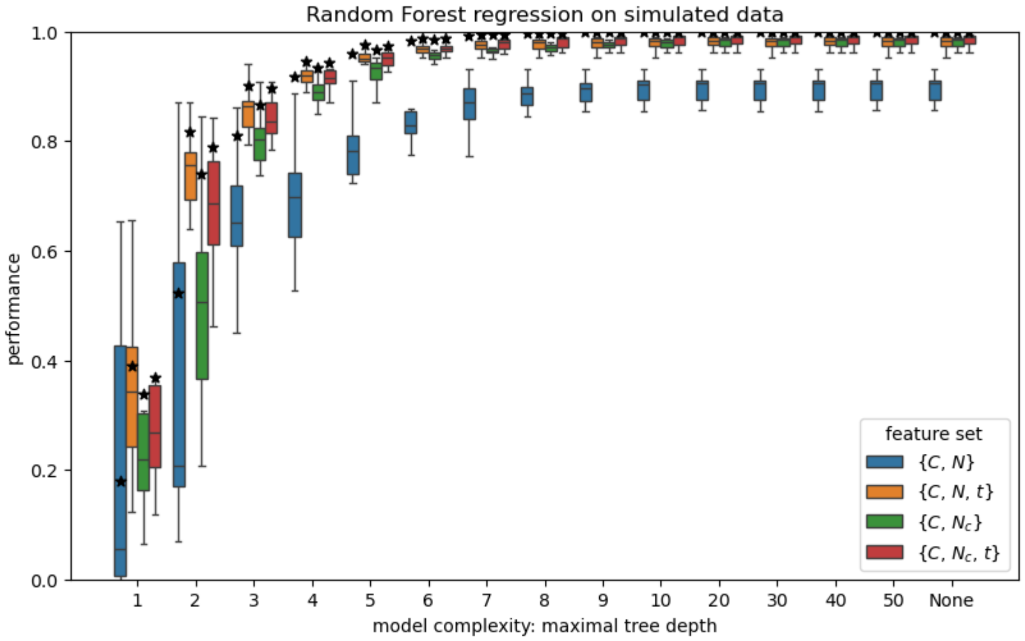

*Figure 2: Black-box Random Forest regression predicting the per-capita growth rate from different feature sets shows that the cumulative population gain predictor $N_c$ outperforms the population size predictor $N$ in simulated data (multiplicative model with five resource dimensions).*

With simulated data, the Random Forest regression qualitatively reproduced the results obtained for experimental data. This shows that the advantage of $N_c$ alone over $N$ alone observed with experimental data does not result from some experimental growth curves being non-monotonic, because we obtain similar results with monotonic, simulated data (figure S2).

#### 2.3 Population size feature importance

Growth data simulated with resource-consumer models intrinsically produces both the overfit described in the main text and its reduction by $N_c$ (figure S3). This does not depend on the underlying resource dimensionality of the resource-consumer models (comparison of models `A` and `M` along the horizontal axis of figure S3).

This does not simply result from $N_c$ reducing, as an integrated quantity, the noise in $N$. If that were the case indeed, our noiseless simulations would not reproduce the result obtained in experiments. But even in the case where noise reduction plays a role, $N_c$ still offers a better indirect access to the latent resource experienced by the growing microbial population compared to $N$. It is similarly known that reporting values of $N_c$ as the area under the growth curve (AUC) allows to reduce the noise observed on measurements of the final experimental value for $N$, namely the yield.

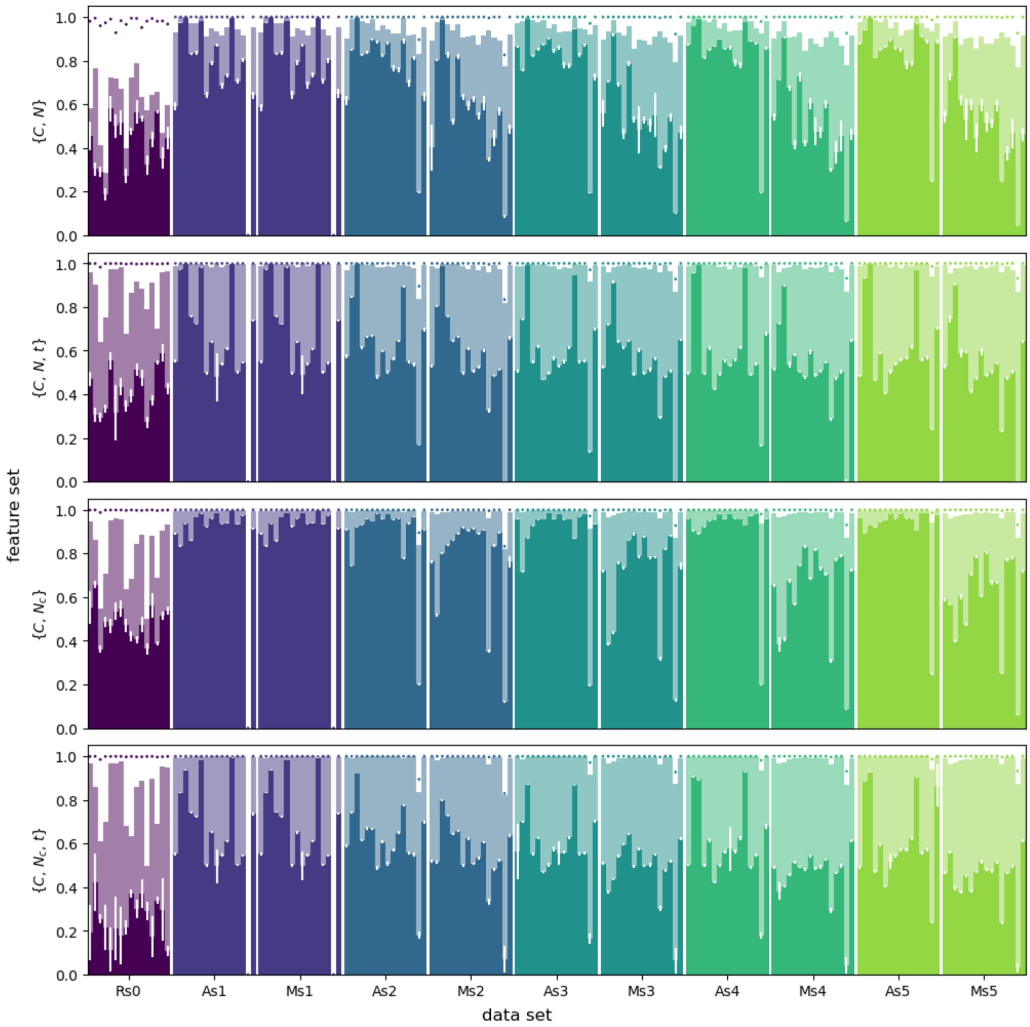

*Figure 3: Performance of a Random Forest regressor, given by the coefficient of determination $R^2$, using different feature sets to predict the per capita growth rate in data obtained: from experiments (`Rs0`); from simulations using an $\alpha$-adjusted model with various levels of resource dimensionality (`Ms1`–`Ms5` and `As1`–`As5`). Each group comprises 16 species and shows, for each, the algorithm performance on the training set (dot), its performance on the testing set (lighter shade), the importance of the population size feature (darker shade), and its standard deviation (white segment).*

### 3 Symbolic regression

#### 3.1 Model search protocol

The algorithm for symbolic regression is explained in detail in Cranmer (2023). In our application, input data comprised the time series of the observed per capita growth rate $\rho_{obs}$ as a predicted variable, together with combinations of the following features: relative culture medium concentration $C$, time $t$, population size $N$, cumulative population gain $N_c$.

For each dataset and each template, the output consisted of a list of symbolic expressions (proposed models) ordered by their parameter number (model complexity). We automatically simplified expressions to reduce artificially inflated complexity values. Note that this complexity value provides an upper bound on a model's parameter number.

We tried to interpret all proposed models in biological terms by focusing on their time-independent part ($\rho$). Without *a priori* excluding other mathematical forms, we paid special attention to models previously reviewed (O'Neill et al., 1989; Zwietering et al., 1990; Ghenu et al., 2024) and to the dynamical Monod model interpreted with our assumption of a constant-rate resource consumption, which corresponds to sums and products of the following mathematical form:

$$\rho = \frac{K_1 K_2 C F^{K_3}}{K_4 + K_2 C F^{K_3}},$$

where $F = \exp(-N_c)$. An example of a biologically interpretable model fitting none of the expected forms appears in figure 5 of the main text (centre right), species `2164`, white triangle annotated as *more complex model* with 8 parameters, which suggested four latent resource dimensions:

$$\frac{1}{N}\frac{dN}{dt} \propto S_1 \times S_2 \times S_3 + S_4.$$

This example illustrates that our search for biologically interpretable models was not limited to expected mathematical forms."

We annotated models manually since no algorithm can reliably determine whether two mathematical expressions are equal in the general case (Richardson, 1969). This manual part had no impact on the pool of automatically proposed models. Due to the arbitrary exponents in the above expression related to the dynamical Monod model, decidability in the case of the corresponding class of models is currently an open question, to our knowledge.

Rules of thumb for the manual part of the model search protocol included the following. The time-dependent adjustment function may be identified and separated from the rest of the proposed growth model whenever that model includes a factor:

$$\alpha = \frac{K}{K + U^m},$$

where $U = \exp(-t)$, and $K$ and $m$ are positive. Then, many models of particular interest could be rearranged into:

$$\frac{\rho}{\alpha} = \frac{\sum_{i'=0}^{l'} K_{i'} C^{i'} F^{i' k_{i'}}}{\sum_{i=0}^{l} K_i C^i F^{i k_i}}.$$

For this form to be analysable as resulting from Monod components, it is necessary that:

- the highest power of $C$ should not be greater in the denominator than in the numerator ($l' \geq l$);
- the denominator should be reducible to units in which $C$ is raised to the power 1, so that the polynomial $\sum_{i=0}^{l} K_i x^i$ should have $l$ real, negative roots.

#### 3.2 Interpretation of model complexity

A few caveats arose concerning the interpretation of model complexity. First, it is often possible to rearrange proposed mathematical expressions so as to minimise the parameter number in a way that we were not able to automatise. We did not go through this highly ineffective task for all models proposed, which means that further analyses relying on parameter numbers imply an additional step of model selection.

Secondly, the expression of certain biologically complex models may in fact reduce to surprisingly few parameters. For instance, five linearly used resources with multiplicative effects on growth (linear $\times$ linear $\times$ linear $\times$ linear $\times$ linear) collapse down to the same number of parameters as the linear-$\times$-linear model, the only difference being the value of the power applied to medium concentration C. Therefore, parameter number should be considered one possible proxy for model complexity, which may or may not perfectly align with biological complexity.

#### 3.3 Simulations for the symbolic regression

The main text abundantly demonstrates the ability of the symbolic regressor to learn a dynamical Monod model relying on a unidimensional resource. Next, we focused on twenty randomly generated simulated cases of a dynamical Monod model relying on a bidimensional resource: ten in which the resources were nonsubstitutable and ten in which they were substitutable. The nonsubstitutable case combined Eq. (1), Eq. (2), Eq. (3), Eq. (4) and Eq. (6), whereas the substitutable case combined Eq. (1), Eq. (2), Eq. (3), Eq. (5) and Eq. (6).

Central parameters were fitted to the experimental data (median across all replicates of all species for each given resource concentration and time point). Random variation was introduced as multiplicative noise affecting each parameter independently, drawn from a lognormal distribution with parameters $\mu = 0$ and $\sigma = 0.2$.

On the one hand, we ran the symbolic regression protocols on data with known resources independently varied. In all cases, the algorithm was able to correctly infer the substitutability of the two resource dimensions (Figure S4). This did not mean that the generative model was inferred exactly. For instance, we could interpret a model proposal as:

$$\frac{1}{N}\frac{dN}{dt} \propto \frac{S_1}{S_1 + K_1} + S_2 \quad \text{instead of generative} \quad \frac{S_1}{S_1 + K_1} + \frac{S_2}{S_2 + K_2},$$

which means that the algorithm:

- correctly inferred two resource dimensions ($S_1$ and $S_2$), which is expected because resources are known in this case;
- correctly inferred their metabolic substitutability (the additive relationship +);
- was unable to infer the nonlinearity in resource $S_2$.

On the other hand, we ran the symbolic regression protocols on data with unknown resources. We focused on the best models in each of the 20 simulated cases, based on $R^2$-loss and based on Akaike's information criterion (AIC). In 8 cases, the form of the generative model was recovered exactly. In 9 other cases, the best model form was a unidimensional approximation of the bidimensional generative model. In 2 cases, a model of the correct dimensionality was proposed but its form did not match exactly that of the generative model. In the remaining case, the best model proposal was more complex. Importantly, all of those groups produced mean and median $R^2$-losses of the same order of magnitude ($10^{-3}$) and mean and median AIC

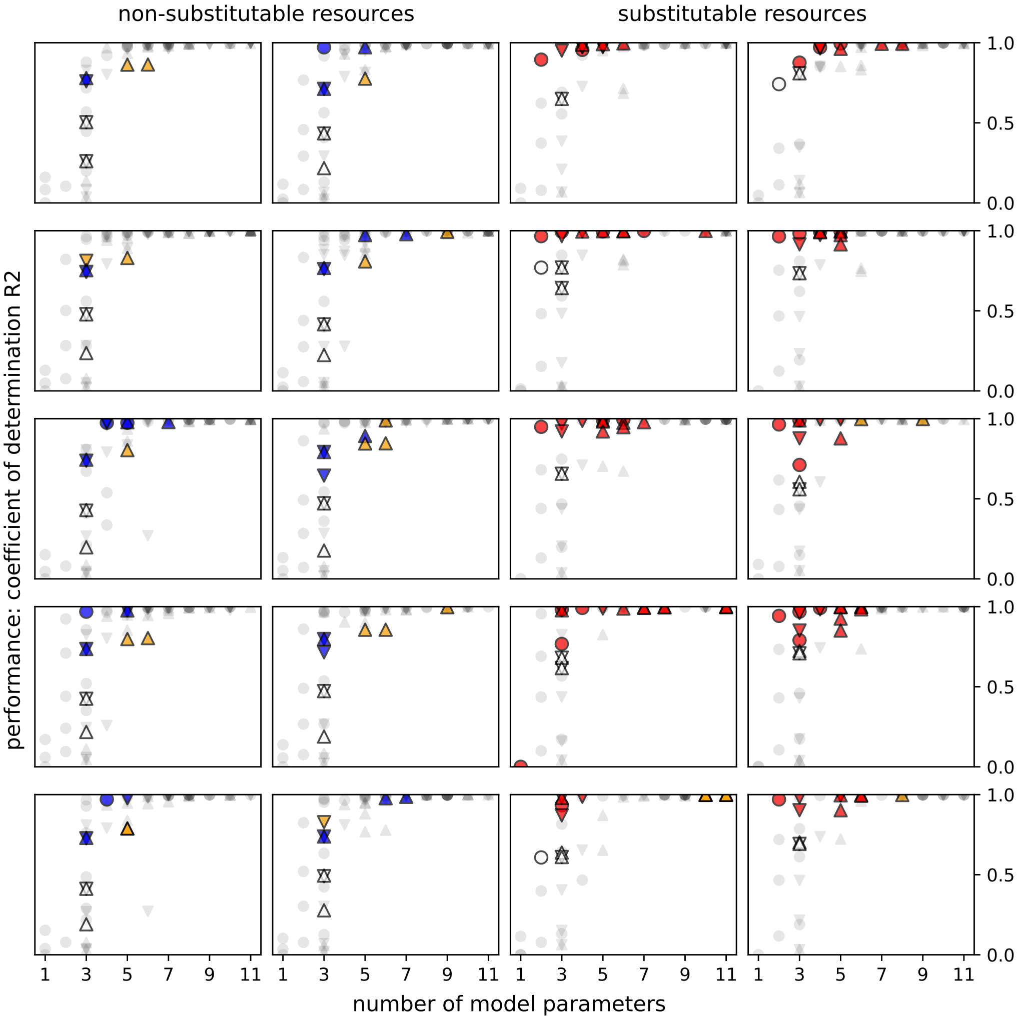

*Figure 4: Fitting performance of models proposed by the symbolic regressor from growth data simulated using a dynamical Monod model relying on a bidimensional resource space, assuming either that the resource dimensions were non-substitutable (left) or substitutable (right). White: unidimensional Monod model; blue: non-substitutable bidimensional Monod model; red: substitutable bidimensional Monod model; orange: higher-dimensional Monod model; gray: not biologically interpretable. Circle: no template; triangle pointing down: Template (7); triangle pointing up: Template (8).*

values of the same order of magnitude ($-10^6$): it was not possible to detect whether the best proposal matched the generative model, without already knowing the generative model.
